# INDELVAR: structure-informed prediction of in-frame indel pathogenicity with calibrated PP3/BP4 thresholds

**DOI:** 10.64898/2026.08.13.737497

**Authors:** Eunhui Ji, Seung Hwan Oh, In-Suk Kim

**Affiliations:** Department of Laboratory Medicine, Pusan National University Yangsan Hospital, Pusan National University School of Medicine, Yangsan, Gyeongsangnam-do, South Korea

**Keywords:** In-frame indels, Variant classification, Protein structure, AlphaFold, Random Forest, Pathogenicity prediction

## Abstract

In-frame insertions and deletions are difficult to interpret because their effects depend on both the sequence change and its protein context. We developed INDELVAR, a random forest model for in-frame insertions and deletions of 1-10 amino acids that integrates 37 features describing AlphaFold-derived wild-type structural context, evolutionary conservation, local sequence change, gene constraint, and curated protein annotations. Pathogenic variants more often affected protein regions with high AlphaFold confidence, low solvent exposure, dense local packing, and strong evolutionary conservation. INDELVAR showed high discrimination in cross-validation with the area under the receiver operating characteristic curve (AUROC) of 0.980, and in an independent test set, an AUROC of 0.977. INDELVAR achieved higher AUROCs than the evaluated methods for both deletions and insertions, although the differences from a recent protein language model-based method were not significant. With separate calibration for deletions and insertions, INDELVAR reached strong evidence on both the pathogenic and benign sides for each type, a range not previously reported for an in-frame indel predictor. In independent testing, all represented evidence intervals met their corresponding likelihood ratio requirements. A precomputed resource provides scores for 372,090 observed in-frame indels mapped to Genome Reference Consortium Human Build 38.

## Background

Small insertions and deletions (indels) in coding regions fall into two types according to their effect on the reading frame. Frameshift indels usually introduce a premature termination codon and are evaluated under the null variant criterion PVS1, which applies when loss of function is an established disease mechanism for the gene and transcript- and position-specific conditions, such as predicted nonsense-mediated decay, are satisfied [1, 2]. In-frame indels change the coding sequence by a multiple of three nucleotides, preserving the reading frame and producing a protein that is intact apart from the inserted or deleted residues [3]. In-frame indels are a frequent cause of inherited disease across diverse genes, but their effects are strongly context-dependent [4, 5]. Because the change leaves the rest of the protein intact, an in-frame indel may be tolerated at one position but abolish function at another [6, 7]. This dependence on sequence and structural context makes in-frame indels hard to interpret, and although they account for approximately 10% of coding ClinVar submissions [8], a disproportionate share is classified as variants of uncertain significance (VUS) [9].

Deep mutagenesis across many proteins identifies loss of fold stability as a major mechanism by which in-frame indels become pathogenic [10]. *CFTR* p.Phe508del destabilizes the nucleotide-binding domain 1 and its interface [11, 12], and similar fold- destabilizing indels occur in *SCN1A* [13], *CRYBA* and *BFSP2* [4], and *CACNA1F* [5]. These effects involve changes to side-chain packing, secondary structure, and hydrogen-bond networks [14]. Deleting or inserting a few residues may be tolerated in a flexible loop but abolish function when it disrupts a buried β-strand or packed core, and this tolerance varies along the protein chain [15].

Most methods that score in-frame indels derive their signal from sequence, evolutionary conservation, or learned sequence representations rather than from explicit three-dimensional protein structure. They span general-purpose predictors such as CADD [16, 17], machine learning models [18], and approaches built on protein language model embeddings [19, 20]. More recently, PON-Del [21] applied AlphaFold-derived features, including secondary structure and solvent accessibility, to in-frame deletions, and structural information improved prediction for this class. These features, however, remain per-residue annotations, much like the secondary structure, disorder, and solvent accessibility that earlier sequence-based predictors already used [22, 23]. AlphaFold now provides structural models for nearly the entire human proteome [24], and structure-aware modeling has already advanced missense variant prediction [25]. What such annotations do not describe is the packing and contact network in which the affected segment sits, and structure-informed prediction of in-frame indels has so far been limited to deletions [21].

Clinical use depends on how a variant is represented and on how a predictor’s score is translated into evidence. Clinical classification follows the American College of Medical Genetics and Genomics/Association for Molecular Pathology (ACMG/AMP) guidelines [1], which weights evidence as Supporting, Moderate, Strong, or Very Strong. The Clinical Genome Resource (ClinGen) Sequence Variant Interpretation Working Group (SVI) placed these strengths on a quantitative scale by expressing each as a likelihood ratio (LR) in a Bayesian model [26, 27]. It then calibrated computational predictors by estimating the local posterior probability of pathogenicity, so that a score maps to a defined evidence strength [28]; several missense predictors now reach strong evidence on this scale [29]. The ClinGen Computational and Variant Classification Working Groups recently extended this calibration to in-frame indels [30]. They evaluated eight prediction tools against ClinVar pathogenic and benign in-frame indels with a Genome Aggregation Database (gnomAD) population reference, applying the same local posterior probability framework to assign score thresholds separately for deletions and insertions. Several of the eight tools reached strong evidence for benignity, whereas none reached strong evidence for pathogenicity; the highest strength for pathogenicity was moderate. The authors recommended choosing an in-frame indel predictor that provides at least moderate pathogenic and supporting benign evidence.

We therefore developed INDELVAR (IN-frame inDEL VARiant pathogenicity predictor) for in-frame insertions and deletions of 1-10 amino acids. INDELVAR integrates 37 features describing wild-type structural context, evolutionary conservation, local sequence change, gene constraint, and curated protein annotations in a random forest model. We then calibrated its scores separately for deletions and insertions to the ACMG/AMP computational evidence criteria PP3 and BP4. We also provide a precomputed resource containing INDELVAR scores and calibrated computational evidence for observed GRCh38 in-frame indels.

## Methods

### Data sources and variant processing

Canonical transcripts were defined using the Matched Annotation from NCBI and EMBL-EBI (MANE) Select v1.5 on Genome Reference Consortium Human Build 38 (GRCh38) [31]. Functional consequence and transcript annotation were generated using the Ensembl Variant Effect Predictor (VEP) 116 [32]. For the training, test, and VUS sets, each variant was represented by the VEP-flagged MANE Select transcript carrying an inframe_deletion or inframe_insertion consequence. Cases with more than one qualifying MANE Select transcript that could not be resolved were excluded.

Gene-level constraint metrics were obtained from gnomAD v4.0 [33]. AlphaFold v6 monomer models were obtained from the AlphaFold Protein Structure Database [34, 35]. Canonical protein sequences were reconstructed from the MANE Select coding sequence. UniProt annotations and AlphaFold-derived structural features were computed only when the reconstructed MANE protein exactly matched a reviewed canonical UniProtKB/Swiss-Prot human sequence from release 2026_02 [36]. When no reviewed accession or exact sequence match was available, the UniProt- and AlphaFold-dependent features were recorded as missing, but the allele was retained.

We selected variants producing an in-frame deletion or insertion of 1-10 amino acids, as determined from the normalized GRCh38 reference and alternate alleles and VEP 116 annotations. We excluded protein-level delins and other complex events. Each allele was applied to the MANE Select coding sequence and translated. We retained variants only when the reconstructed mutant protein agreed with VEP’s protein-level annotation.

### Training and test sets

Clinical variants were taken from the ClinVar releases of 6 June 2024 and 4 June 2026 [8]. We required a review status of at least one star. The pathogenic/likely pathogenic (P/LP) class comprised Pathogenic, Likely pathogenic, and Pathogenic/Likely pathogenic records; the benign/likely benign (B/LB) class comprised Benign, Likely benign, and Benign/Likely benign records. Records with conflicting, mixed, or non-definitive classifications were excluded, and duplicate alleles were collapsed after normalization. Uncertain significance (VUS) entries from the 2026 set were reserved for a separate VUS analysis after the model and calibration thresholds had been fixed.

The training set comprised all eligible P/LP and B/LB alleles from the 2024 ClinVar (n = 4,927: 2,103 P/LP and 2,824 B/LB; 3,510 deletions and 1,417 insertions). The test set comprised eligible P/LP and B/LB alleles from the 2026 ClinVar after excluding the training set overlap by exact normalized allele. The resulting test set contained 1,262 alleles (764 P/LP and 498 B/LB; 940 deletions and 322 insertions).

### Feature design

INDELVAR integrates 37 features in five groups: protein structure (n = 11), evolutionary conservation (n = 2), sequence context (n = 8), gene constraint (n = 4), and curated protein annotations (n = 12) (**Fig. 1**). **Supplementary Table S1** lists each feature’s definition, calculation, source, and handling of missing data.

**Figure 1.**
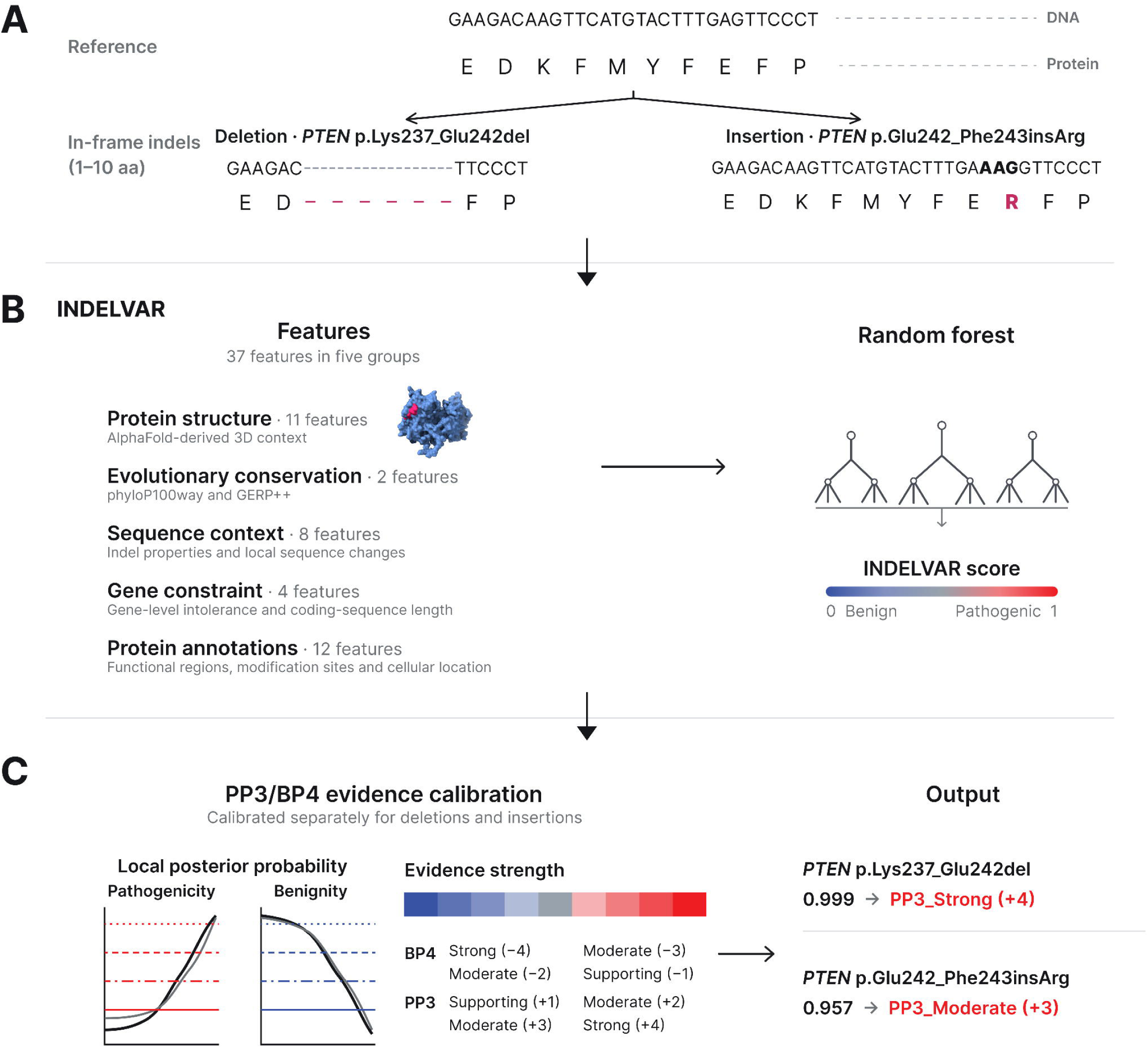
Overview of INDELVAR. (A) Examples of a normalized GRCh38 in-frame deletion and insertion within the supported range of 1-10 amino acids. (B) Each variant is represented by 37 features in five groups: protein structure (n = 11), evolutionary conservation (n = 2), sequence context (n = 8), gene constraint (n = 4), and protein annotations (n = 12). The AlphaFold structure of PTEN illustrates the structural context of the deleted residues. A 1,000-tree random forest combines the features into an INDELVAR score ranging from 0 to 1. (C) Scores are mapped to PP3 or BP4 evidence strengths using thresholds calibrated separately for deletions and insertions. *PTEN* p.Lys237_Glu242del received a score of 0.999, corresponding to PP3 strong evidence (+4), whereas *PTEN* p.Glu242_Phe243insArg received a score of 0.957, corresponding to PP3 moderate evidence (+3).

*Protein structure* (n = 11). Indel effects depend on local structural context [10]. We calculated 11 features from the wild-type AlphaFold model to describe the structural context at each variant site. Structural features were calculated from the wild-type AlphaFold model over the deleted residues for deletions and the four wild-type residues surrounding the insertion junction for insertions. These comprised mean predicted local distance difference test (pLDDT), helix and sheet fractions, relative solvent accessibility (RSA), weighted contact number (WCN) [37], contact density [14], bridging contacts [38], secondary-structure-element break and fraction affected, hydrogen-bond density, buried hydrophobic fraction [39]. PyDSSP 0.9.1 provided the three-state secondary-structure assignments and hydrogen-bond map based on the dictionary of secondary structure of proteins (DSSP) framework [40], and RSA was calculated with FreeSASA 2.2.1 [41] and normalized by the residue-specific theoretical maximum solvent-accessible surface areas reported by Tien et al. [42]. These features describe the wild-type structural context and do not model the mutant protein structure.

*Evolutionary conservation* (n = 2). Because evolutionarily conserved regions are generally less tolerant of sequence changes [43], we calculated mean phyloP100way [44] and GERP++ [45] scores over the deleted reference bases for deletions and the 12 coding bases corresponding to the four wild-type junction residues for insertions.

*Sequence context* (n = 8). Indel tolerance differs in low-complexity and repetitive regions [46], and exonic variants close to splice junctions may disrupt splicing [47]. The sequence-context features comprised indel size, size relative to protein length, indel type, local amino-acid entropy, distance to the nearest internal exon boundary, and variant-induced changes in local hydropathy, entropy, and charge. Local amino-acid entropy was calculated as Shannon entropy [48] over the variant window expanded by up to 10 residues on each side. Changes in local hydropathy, entropy, and charge were defined as the mutant value minus the wild-type value; hydropathy and charge were calculated using the Kyte-Doolittle hydropathy scale [49] and the net-charge scale [50], respectively. Wild-type and mutant local sequences were reconstructed from translated MANE Select coding sequences using up to 10 flanking residues on each side.

*Gene constraint* (n = 4). Gene-level intolerance to protein-altering variation provides information complementary to variant-level features [51]. We included probability of loss-of-function intolerance (pLI), loss-of-function observed/expected upper-bound fraction (LOEUF), and the missense constraint Z-score from gnomAD v4.0 [33], together with the MANE Select coding sequence length excluding the stop codon [31].

*Protein annotations* (n = 12). Indels overlapping functional protein elements are more likely to impair protein function [52]. Five local features derived from UniProt [36] described the number of overlapping annotated regions, overlap with a repeat, the number of nearby post-translational modification sites, proximity to a disulfide-bonded cysteine, and overlap with an active or binding site. Functional and post-translational modification sites have previously been associated with pathogenic variation [53, 54]. Six multi-label indicators represented nuclear, cytoplasmic, membrane, mitochondrial, secreted, and other subcellular locations. A seventh indicator identified reviewed UniProt records without a structured subcellular-location annotation.

Allele frequency was excluded because it is evaluated separately under the ACMG/AMP population-frequency criteria, and including it could double-count population evidence when PP3 or BP4 is applied [28]. Missing numeric values were imputed using the median from the training portion of each cross-validation fold and, for the final model, the median from the full training set.

### Model development and assessment

Random forest, XGBoost [55], and elastic-net logistic regression [56] were compared in the training set by event-grouped nested cross-validation, using fourfold inner cross-validation within five outer folds and the mean AUROC across indel types for model selection. We evaluated complete grids of 480 random-forest and 700 elastic-net configurations, together with 480 prespecified configurations from a 61,440-combination XGBoost search space; for the random forest, mtry, terminal-node size, sampling fraction, and maximum-node limit were tuned, whereas the number of trees was fixed at 1,000 and assessed separately for convergence. The selected random forest was fitted using randomForest version 4.7-1.2 [57, 58] with mtry = 7, nodesize = 1, sampling fraction = 1.0 with replacement, and no maximum-node limit (S**upplementary Fig. S1**). The INDELVAR score was defined as the proportion of trees voting for the P/LP class, ranging from 0 to 1.

Internal performance was assessed from the pooled out-of-fold predictions of the five outer folds. Variants from the same transcript that produced identical mutant protein sequences were assigned to the same fold. Imputation medians were estimated separately within each analysis fold. Each training variant received one out-of-fold prediction, and the pooled out-of-fold predictions were used to assess internal discrimination and for subsequent evidence calibration.

The final model was fitted to the complete training set using the same settings and was then applied to the independent test set. Preprocessing parameters were derived from the complete training set, and no model refitting, recalibration, or modification of the preprocessing procedure was performed using the test data. Discrimination was evaluated using the area under the receiver operating characteristic curve (AUROC), both overall and separately for deletions and insertions. Confidence intervals (CIs) were estimated using 2,000 percentile bootstrap replicates stratified by variant class and indel type. Continuous INDELVAR scores were used throughout, and no decision threshold was selected using the test set.

Feature importance was quantified using the mean decrease in Gini impurity and normalized as a percentage of the total importance across all 37 features. Pairwise associations among model features were also evaluated in the training set using pairwise complete observations and measures appropriate to the corresponding variable types.

### Comparison with other prediction methods

We compared INDELVAR with CADD v1.7 [16], FATHMM-indel [59], MutPred-Indel [18], PON-Del [21], and the IndeLLM Siamese model [20], together with phyloP100way and GERP++ as conservation baselines. Comparison methods were selected based on their relevance to in-frame indel interpretation, methodological diversity, and accessibility for reproducible large-scale evaluation. Methods that incorporate allele frequency in their predictions were omitted to avoid double-counting population evidence [28]. We also considered the availability of training data to assess overlap with the INDELVAR test set where possible. PON-Del was evaluated for deletions only.

Deletions and insertions were evaluated separately using each method’s continuous score in its prespecified direction. Analyses were restricted to alleles for which the corresponding score was available. Comparisons with INDELVAR were performed on the same alleles scored by both methods. Differences in AUROC were assessed using paired DeLong tests, with Holm correction for multiple comparisons. CIs for each method’s AUROC and for paired AUROC differences were estimated using 2,000 bootstrap replicates stratified by variant class and indel type.

To address potential type 1 circularity [60], overlap between comparator training data and the INDELVAR test set was assessed where possible. Overlap was defined using exact normalized genomic alleles when available. For IndeLLM, whose released training data did not contain complete genomic allele information, overlap was assessed at the protein-event level. Comparator versions and access routes are provided in **Supplementary Table S2**.

### ACMG/AMP evidence calibration

INDELVAR scores were calibrated to ACMG/AMP computational evidence (PP3 and BP4) strengths using the local posterior probability method of Pejaver et al. [28], applied separately to deletions and insertions following Abderrazzaq et al. [30]. Calibration used the complete event-grouped out-of-fold predictions for 4,927 training variants. A population reference comprising 23,253 in-scope variants from gnomAD—16,408 deletions and 6,845 insertions—was scored using the final model and used to assess local support. Local posterior probabilities were estimated within adaptive score windows that were expanded until they contained sufficient calibration data, defined as at least 100 ClinVar variants and 3% of the corresponding indel-type-specific gnomAD reference set.

The prior probabilities of pathogenicity were fixed at 4.6% for deletions and 0.8% for insertions [30]. Target likelihood ratios for evidence points +1 to +4 were 2.39, 5.69, 13.59, and 32.42 for deletions and 3.25, 10.55, 34.28, and 111.34 for insertions; reciprocal targets were used for the corresponding BP4 evidence points. Thresholds were derived from the continuous INDELVAR score using 10,000 bootstrap resamples and the one-sided 95% discounting procedure described by Pejaver et al. [28]. A calibration target was considered unreached when more than 5% of bootstrap resamples produced no valid threshold.

The resulting deletion- and insertion-specific thresholds were applied unchanged to the independent test set. For each PP3 and BP4 evidence tier, the observed likelihood ratio was compared with the corresponding target, and 95% CIs were estimated using 2,000 bootstrap replicates stratified by variant class where estimable.

### ClinVar VUS analysis

The VUS analysis set consisted of 20,639 variants from the 2026 ClinVar release: 14,473 deletions and 6,166 insertions. All met the study eligibility criteria, and none were included in the training or test sets.

Variants were assigned to PP3 or BP4 evidence tiers using the calibrated deletion- and insertion-specific thresholds. Variants that did not meet any threshold remained computationally indeterminate. The assigned tiers represent computational evidence only and were not treated as variant reclassification.

### Precomputed score resource

To construct the INDELVAR resource, we scored eligible GRCh38 in-frame indels compiled from the ClinVar releases of 6 June 2024 and 4 June 2026 [8], the International Genome Sample Resource (IGSR)/1000 Genomes 20220422 panel [61], and dbSNP [62]. Because MANE Select is defined per gene, variants affecting MANE Select transcripts from multiple genes were represented and scored separately for each transcript.

### Statistical analysis

Statistical analyses were performed in R 4.5.3. Continuous and ordinal features were compared between P/LP and B/LB variants separately for deletions and insertions using two-sided Mann-Whitney U tests. Feature comparisons were adjusted for multiple testing using the Holm method. Associations among model features were calculated from pairwise complete observations using measures selected according to variable type. Adjusted *P* < 0.05 was considered statistically significant.

## Results

### Pathogenic in-frame indels preferentially occur in ordered and buried protein regions

To assess the biological features that distinguish P/LP from B/LB in-frame indels, we compared structural features between the two classes, analyzing deletions and insertions separately. Structural annotations were available for 4,273 of the 4,927 training variants, comprising 3,070 deletions and 1,203 insertions. For both indel types, P/LP variants occurred in regions with higher AlphaFold pLDDT, lower RSA, and greater contact density than B/LB variants (adjusted *P* < 0.001 for all comparisons; **Fig. 2A-C**). P/LP variants also had higher local helix fractions than B/LB variants for both deletions and insertions (adjusted *P* < 0.001; **Fig. 2D**).

**Figure 2.**
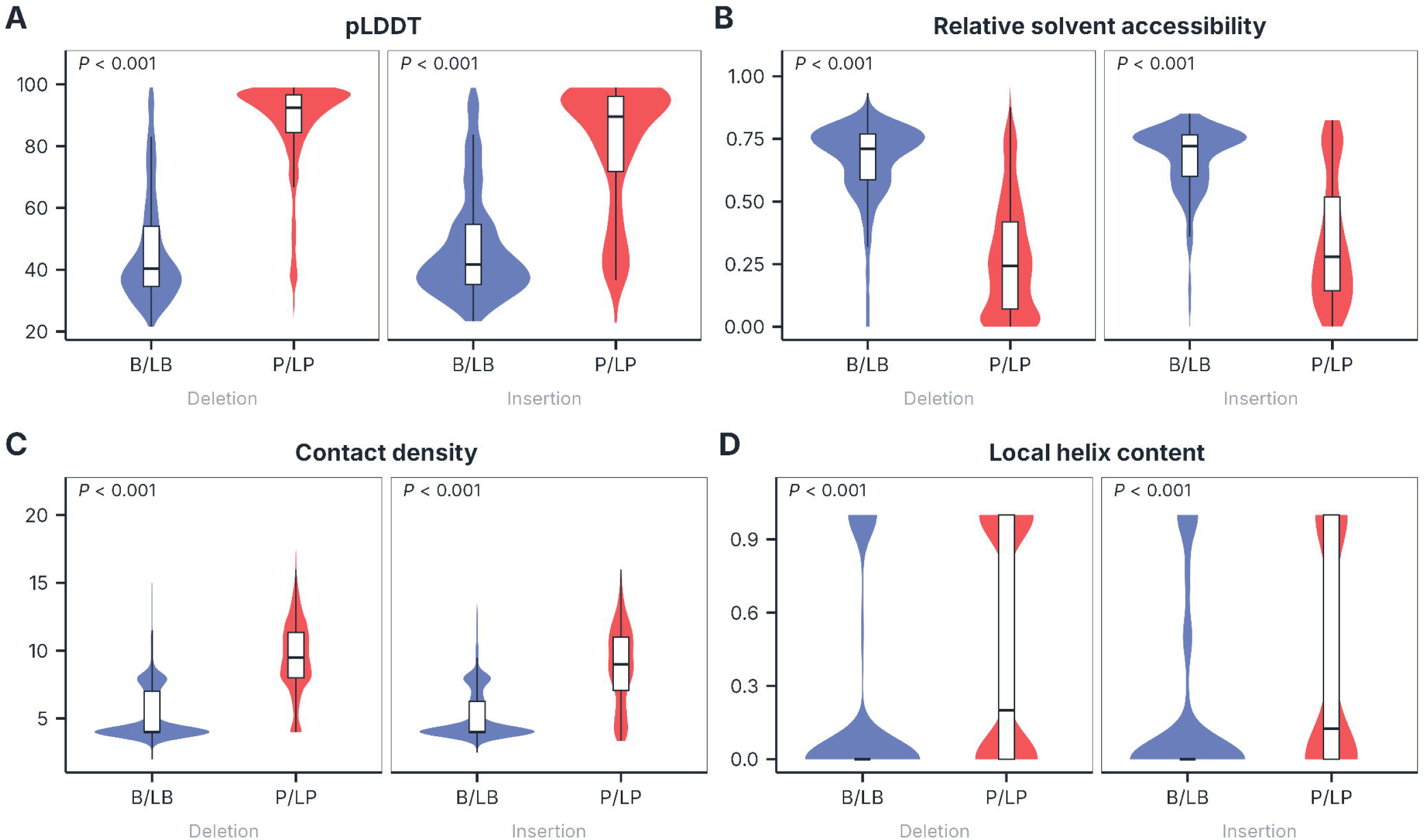
Structural features associated with in-frame indel pathogenicity. Raw feature values from the training set are shown before imputation, with insertions and deletions analyzed separately. Variants without structural annotations were excluded, leaving 1,203 insertions and 3,070 deletions. (A) AlphaFold predicted local distance difference test (pLDDT) and (B) relative solvent accessibility (RSA) in benign/likely benign (B/LB) and pathogenic/likely pathogenic (P/LP) variants. Violin plots show the distributions, with embedded box plots indicating the median and interquartile range. (C) Contact-density distributions in B/LB and P/LP variants, shown separately for insertions and deletions. (D) Distribution of the fraction of residues classified as helix within each variant window in B/LB and P/LP variants. Two-sided Mann-Whitney U tests were used for all four panels. *P* values were adjusted using the Holm method.

### Pathogenic and benign in-frame indels differ in conservation, length, and local sequence change

We next examined whether pathogenicity was associated with evolutionary conservation, the extent of the protein change, and its effects on local hydropathy and sequence entropy. Conservation scores were available for 4,919 training variants, whereas analyses of indel length and changes in local hydropathy and sequence entropy included all 4,927 training variants. P/LP variants occurred at substantially more conserved positions than B/LB variants for both deletions and insertions. Median phyloP100way scores were 4.51 versus 1.06 for deletions and 4.05 versus 0.95 for insertions (adjusted *P* < 0.001; **Fig. 3A**).

**Figure 3.**
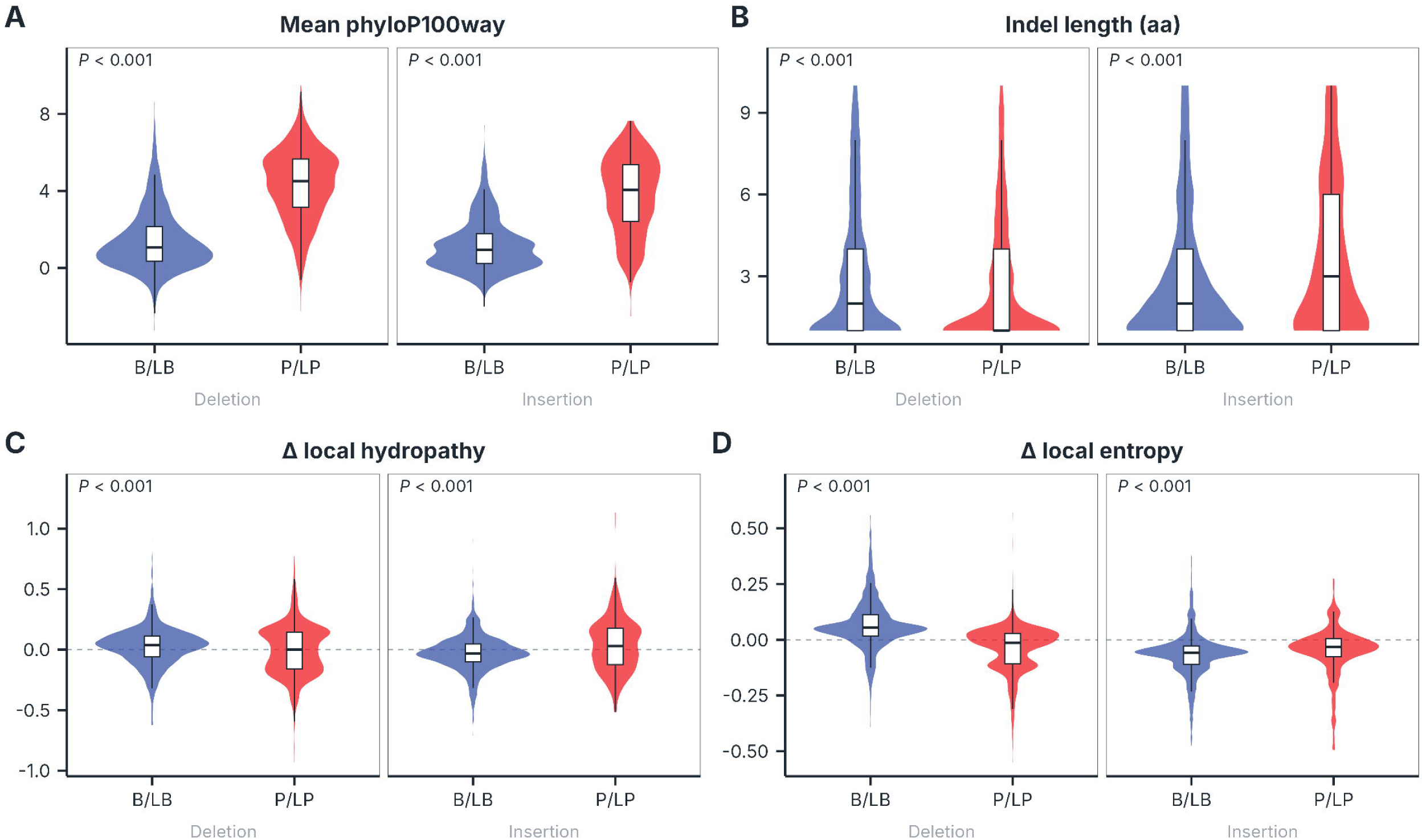
Evolutionary and local sequence features associated with in-frame indel pathogenicity. Raw feature values from the training set are shown before imputation, with insertions and deletions analyzed separately. Complete cases included 1,409 insertions and 3,510 deletions for panel A and 1,417 insertions and 3,510 deletions for panels B-D. (A) Mean phyloP100way conservation score at the affected positions. (B) Indel length in amino acids. (C) Change in local hydropathy and (D) change in local sequence entropy. Changes were calculated as the mutant value minus the wild-type value; dashed lines indicate no change. Violin plots show the distributions, with embedded box plots indicating the median and interquartile range. Two-sided Mann-Whitney U tests were used for all panels. Displayed *P* values were adjusted using the Holm method.

Indel length showed different patterns for deletions and insertions. P/LP deletions were slightly shorter than B/LB deletions, whereas P/LP insertions were longer than B/LB insertions (adjusted *P* < 0.001; **Fig. 3B**). Mutant-minus-wild-type changes in local hydropathy and sequence entropy also differed between P/LP and B/LB variants, with opposite directions for deletions and insertions (adjusted *P* < 0.001; **Fig. 3C-D**). P/LP deletions showed lower changes in both properties, whereas P/LP insertions showed higher changes than their B/LB counterparts.

### INDELVAR distinguishes pathogenic from benign in-frame indels in both cross-validation and independent testing

Together, these findings highlighted structural context, evolutionary conservation, and local sequence change as informative dimensions of in-frame indel pathogenicity. We combined these properties with gene constraint and curated protein annotations into a 37- feature model. In event-grouped nested cross-validation, random forest was selected, with deletion and insertion AUROCs of 0.9772 and 0.9833, respectively, from pooled out-of-fold predictions across the outer folds (mean AUROC across indel types, 0.9802). The corresponding mean AUROCs were 0.9789 for XGBoost and 0.9541 for elastic-net logistic regression (**Supplementary Fig. S1**). Random forest was used for the final INDELVAR model. The five highest-ranked features by mean decrease in Gini impurity were pLDDT, local sequence entropy, phyloP100way, RSA, and contact density (**Fig. 4A**). The complete feature ranking is provided in **Supplementary Fig. S2**, and pairwise associations among the 37 model features are summarized in **Supplementary Fig. S3**.

**Figure 4.**
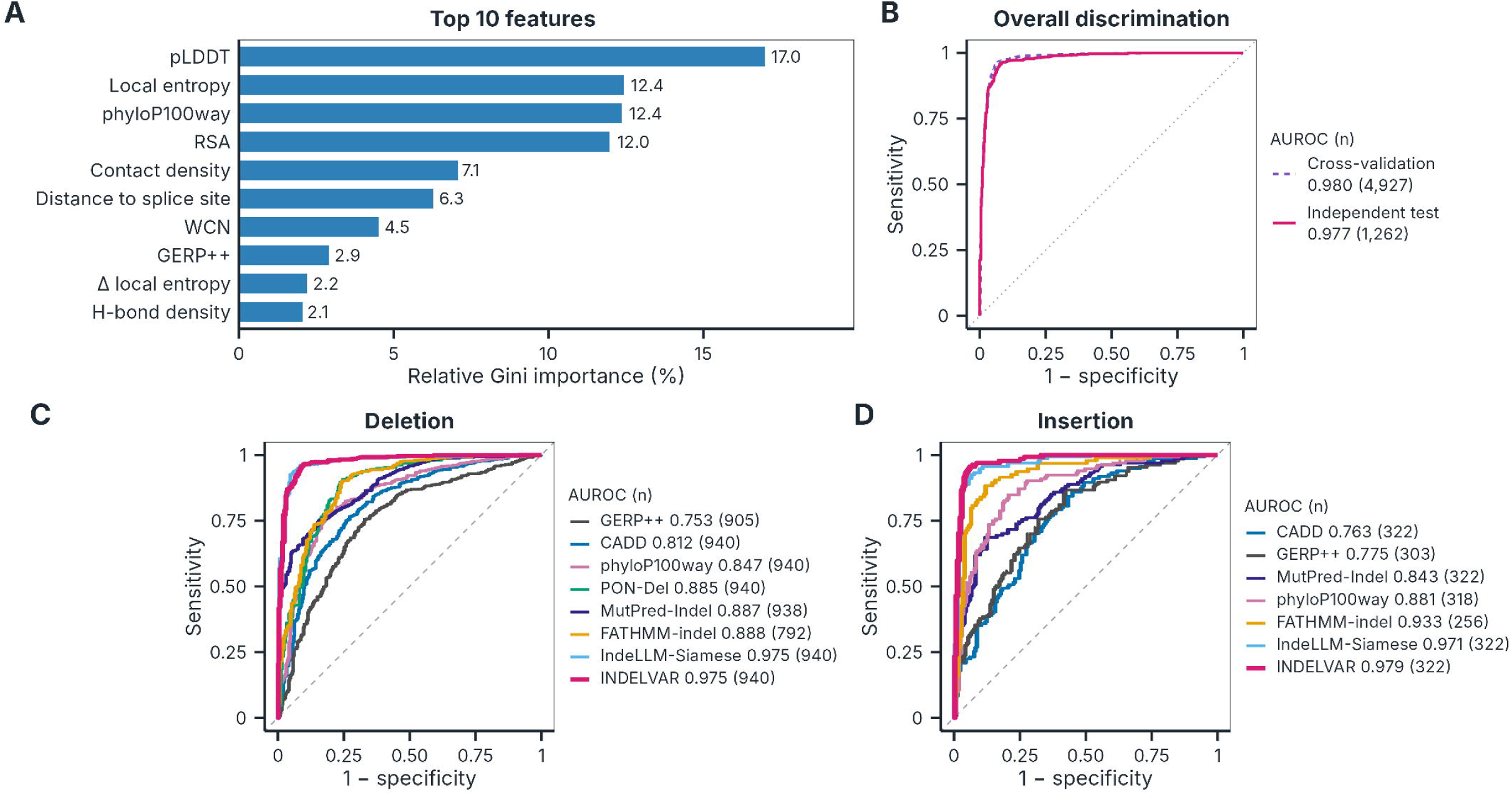
Development and assessment of INDELVAR. (A) Relative Gini importance of the ten highest-ranked features in the final random forest, expressed as a percentage of the total mean decrease in Gini impurity. The complete ranking of all 37 features, colored by feature group, is shown in Supplementary Figure S2. Importance reflects feature use within the fitted forest and should not be interpreted as a causal effect. (B) Receiver operating characteristic curves for pooled out-of-fold predictions from the five outer folds of event-grouped nested cross-validation (n = 4,927; AUROC, 0.980; 95% CI, 0.976-0.983) and predictions in the independent temporal test set (n = 1,262; AUROC, 0.977; 95% CI, 0.968-0.984), overlaid in a single panel. The legend reports the AUROC for each curve. Variants from the same transcript that produced identical mutant protein sequences were kept in the same cross-validation fold. (C, D) Receiver operating characteristic curves for INDELVAR and the comparison methods in the deletion (n = 940; C) and insertion (n = 322; D) test subsets. Each comparison method was evaluated on the variants for which its score was available; the corresponding sample size and AUROC are shown in each panel’s legend. PON-Del was available only for deletions. Formal comparisons with INDELVAR used paired DeLong tests with Holm correction and were restricted to variants scored by both methods. CIs for AUROC and paired differences used 2,000 bootstrap replicates stratified by variant class and indel type.

In pooled out-of-fold predictions for the training set (n = 4,927), INDELVAR achieved an AUROC of 0.980 (95% CI, 0.976-0.983). On the independent test set (n = 1,262), INDELVAR achieved an AUROC of 0.977 (95% CI, 0.968-0.984; **Fig. 4B**).

### INDELVAR achieves the highest AUROCs for both deletions and insertions

INDELVAR had the highest AUROC among the evaluated methods for both deletions and insertions (**Fig. 4C, D**), although the differences from the closest-performing method were not statistically significant.

For deletions, INDELVAR achieved an AUROC of 0.975. A paired comparison with the IndeLLM Siamese model showed no significant difference (AUROC 0.975; Holm-adjusted *P* = 1.000). INDELVAR showed higher discrimination than FATHMM-indel, MutPred-Indel, PON-Del, CADD, phyloP100way, and GERP++, which achieved AUROCs of 0.888, 0.887, 0.885, 0.812, 0.847, and 0.753, respectively (all adjusted *P* < 0.001).

For insertions, INDELVAR also achieved a higher AUROC than the IndeLLM Siamese model (0.979 vs 0.971), although the difference was not significant (Holm-adjusted *P* = 0.179). INDELVAR also exceeded FATHMM-indel, MutPred-Indel, CADD, phyloP100way, and GERP++, which yielded AUROCs of 0.933, 0.843, 0.763, 0.881, and 0.775, respectively (all adjusted *P* < 0.05).

### INDELVAR reaches strong evidence of pathogenicity and benignity for both indel types upon calibration

Calibration yielded thresholds for supporting, moderate, and strong evidence on both the pathogenic and benign sides for deletions and insertions (**Fig. 5A-B**; **Table 1**). Score thresholds were determined from the final local posterior probability curves generated separately for deletions and insertions. For insertions, the −3 and −4 benign thresholds coincided at 0.014, so no separate −3 score interval was defined. The resulting 15 distinct evidence intervals were applied to the independent test set.

**Figure 5.**
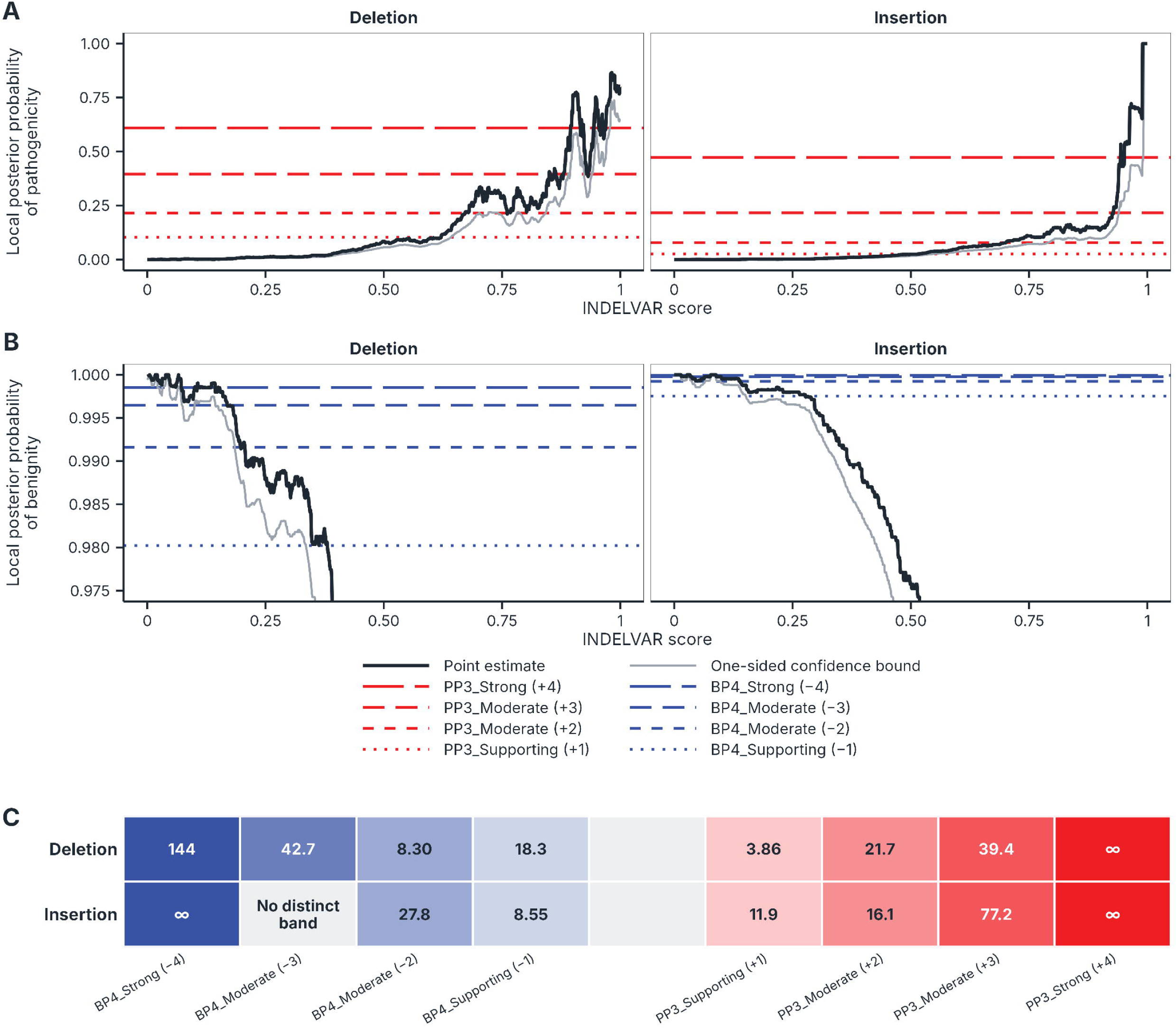
Calibration and independent test-set evaluation of INDELVAR evidence thresholds. Local posterior probabilities were calculated separately for deletions and insertions using indel-type-specific pathogenicity priors. (A) Local posterior probability of pathogenicity, with red horizontal lines, and (B) local posterior probability of benignity, with blue horizontal lines; each panel shows deletions on the left and insertions on the right. The horizontal lines represent the posterior probability thresholds for supporting, moderate, and strong evidence. The black curves represent the posterior probability estimated from the event-grouped nested cross-validation out-of-fold predictions of the 4,927 training variants. The grey curves represent one-sided 95% confidence bounds calculated from 10,000 bootstrap samples of that set, in the direction of more stringent thresholds. The points at which the grey curves intersect the horizontal lines represent the thresholds for the relevant intervals. (C) Observed likelihood ratios within the calibrated score intervals in the independent test set. Each cell shows the observed likelihood ratio; the number of variants in each interval is given in Supplementary Table S3. Infinite values indicate intervals containing no incorrect predictions. The insertion −3 cutoff coincided with the −4 cutoff and therefore had no distinct interval. Every other interval was represented in the test set and met the likelihood ratio requirement for its assigned evidence strength.

**Table 1.**
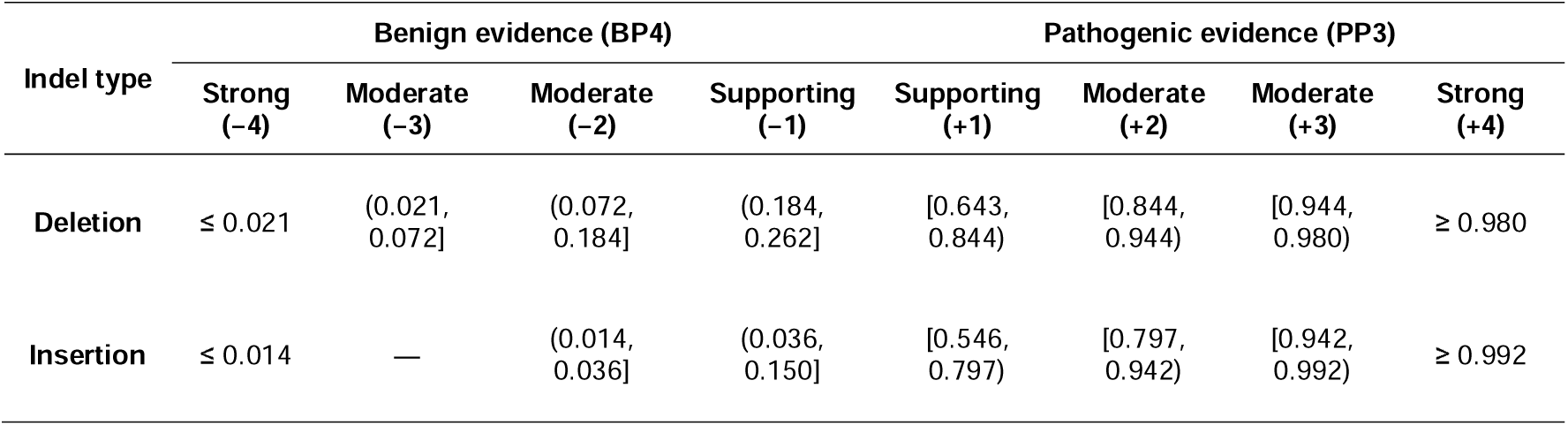
INDELVAR score intervals corresponding to PP3 and BP4 evidence strengths for in-frame deletions and insertions. Higher INDELVAR scores indicate stronger evidence for pathogenicity. Calibration was performed separately for deletions and insertions. For insertions, the BP4 moderate (−3) cutoff coincided with the strong (−4) cutoff, leaving no distinct −3 interval. Scores within 0.262 < score < 0.643 for deletions and 0.150 < score < 0.546 for insertions receive neither PP3 nor BP4 evidence.

| Indel type | Benign evidence (BP4) |  |  |  | Pathogenic evidence (PP3) |  |  |  |
| --- | --- | --- | --- | --- | --- | --- | --- | --- |
|  | Strong (−4) | Moderate (−3) | Moderate (−2) | Supporting (−1) | Supporting (+1) | Moderate (+2) | Moderate (+3) | Strong (+4) |
| **Deletion** | $\leq 0.021$ | (0.021, 0.072] | (0.072, 0.184] | (0.184, 0.262] | [0.643, 0.844) | [0.844, 0.944) | [0.944, 0.980) | $\geq 0.980$ |
| **Insertion** | $\leq 0.014$ | — | (0.014, 0.036] | (0.036, 0.150] | [0.546, 0.797) | [0.797, 0.942) | [0.942, 0.992) | $\geq 0.992$ |

To evaluate the calibrated thresholds on unseen variants, likelihood ratios were estimated for the evidence intervals represented in the independent test set. Each evaluable interval met the likelihood ratio requirement for its assigned evidence strength (**Fig. 5C**; **Supplementary Table S3**). The deletion +4 interval (n = 126), the insertion +4 interval (n = 5), and the insertion −4 interval (n = 87) contained no incorrect predictions, resulting in infinite likelihood ratios. All 15 distinct intervals were represented in the test set and met their respective requirements.

### INDELVAR assigns PP3 or BP4 evidence to most ClinVar in-frame indel VUS

The calibrated thresholds were applied to 20,639 ClinVar VUS, comprising 14,473 deletions and 6,166 insertions. Overall, 16,813 variants (81.5%) received computational evidence: 8,710 (42.2%) received PP3 evidence and 8,103 (39.3%) received BP4 evidence, while 3,826 (18.5%) remained computationally indeterminate (**Fig. 6**). Among deletions, 44.3% received PP3 evidence and 37.7% received BP4 evidence, compared with 37.3% and 42.9% of insertions, respectively. The remaining 18.0% of deletions and 19.7% of insertions were computationally indeterminate. The insertion BP4 −3 tier was not assignable because its calibrated cutoff coincided with BP4 −4, leaving no distinct score interval.

**Figure 6.**
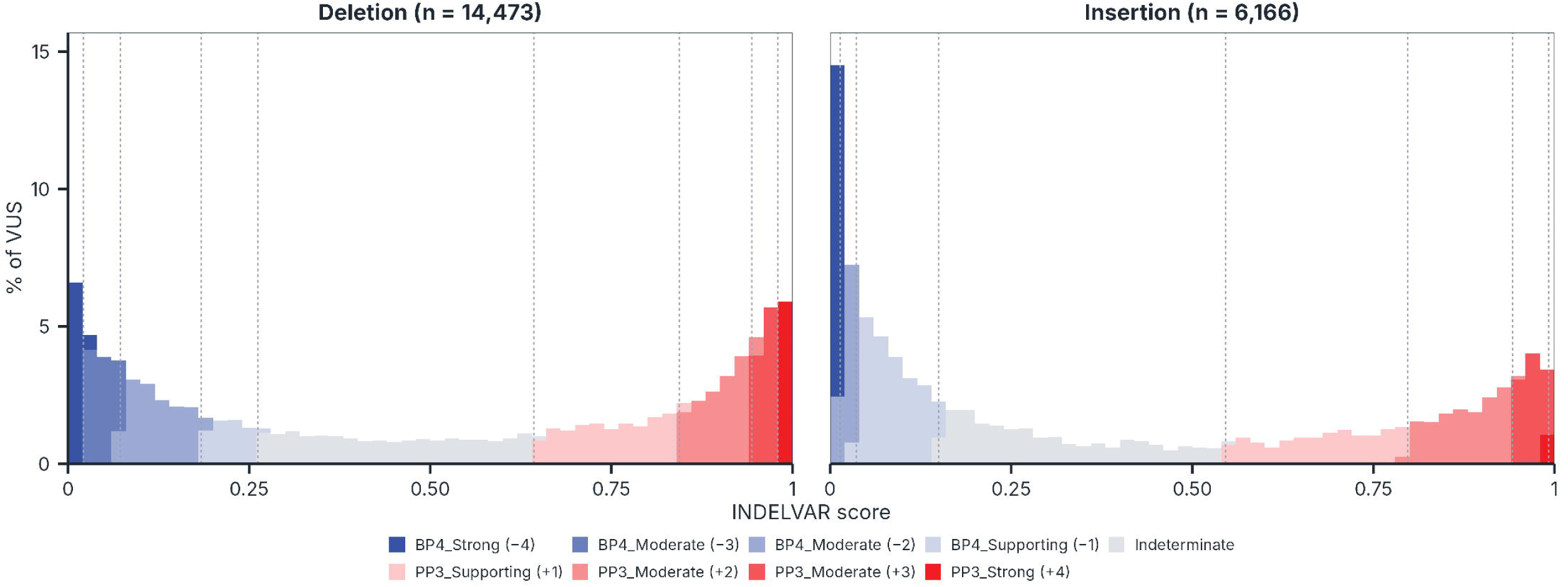
Computational evidence assigned to ClinVar variants of uncertain significance. Distribution of INDELVAR scores among 20,639 in-frame indel variants of uncertain significance in the ClinVar 2026 analysis set. Deletions and insertions are shown separately, with percentages calculated within each panel. Colors indicate the strongest PP3 or BP4 tier reached, and dashed lines show the indel-type-specific calibrated cutoffs. The insertion BP4 −3 tier has no distinct interval because its cutoff coincides with BP4 −4. Assignments represent computational evidence, not clinical reclassification. Bin width, 0.02.

### INDELVAR provides precomputed scores for 372,090 observed in-frame indels

The precomputed resource contained 372,518 records representing 372,090 distinct GRCh38 in-frame indels across 18,805 genes. 215 variants affected the MANE Select transcripts of more than one gene and were represented by separate records for each transcript. The resource comprised 247,629 deletion and 124,889 insertion records. Among all records, 133,564 (35.9%) received PP3 evidence and 165,285 (44.4%) received BP4 evidence, while 73,669 (19.8%) were computationally indeterminate. Each record reports the 37 raw model features, the INDELVAR score, and the corresponding computational evidence, in 60 columns.

## Discussion

INDELVAR was developed on the premise that the effect of an in-frame indel depends not only on the sequence change but also on where the affected residues lie within the folded protein. The preferential occurrence of pathogenic variants at high-confidence, buried, densely packed, and evolutionarily conserved sites supports the relevance of this context. INDELVAR combined AlphaFold-derived structural features with sequence and gene-level information and showed high discrimination in both cross-validation and independent testing. Its AUROCs were the highest among the evaluated methods for both deletions and insertions, although the differences from a recent protein language model-based method were not statistically significant. Upon calibration, INDELVAR also reached strong evidence of both pathogenicity and benignity for deletions and insertions.

Abderrazzaq et al. calibrated eight in-frame indel predictors and found that none of them met the evidence-strength standard previously proposed for missense predictors—strong evidence on the pathogenic side and moderate evidence on the benign side [28, 30]. They therefore recommended selecting an in-frame indel predictor that reaches at least moderate evidence of pathogenicity and supporting evidence of benignity for both deletions and insertions. INDELVAR exceeds this proposed minimum and meets the earlier missense-based standard for both indel types. To our knowledge, INDELVAR is the first calibrated in-frame indel predictor to meet this standard for both deletions and insertions. Because the studies used different variant sets, this comparison concerns calibrated evidence strength rather than head-to-head predictive performance. All 15 distinct evidence intervals were represented in the independent test set and met the likelihood ratio requirement for their assigned strength. The insertion +4 interval contained only five test variants, so further evaluation in larger independent datasets will be needed.

The structural signal captured by INDELVAR is consistent with prior evolutionary and experimental evidence. Indels are depleted from structured protein regions [63], and deep indel mutagenesis has shown lower tolerance in β-strands than in helices [10]. Experimental deletion mapping of enhanced green fluorescent protein also identified WCN, a measure of local packing density, as the strongest single structural predictor of deletion tolerance [64]. In a buried and densely packed environment, even a short length change may disrupt core packing, hydrogen-bond networks, or secondary-structure continuity; such mechanisms have been described for *CFTR* p.Phe508del [11, 12], *SCN1A* p.Met1841del [13], and *BFSP2* p.Glu233del [4]. The greater conservation observed at pathogenic sites is likewise consistent with purifying selection against damaging in-frame indels [63]. These findings support explicit representation of the local wild-type three-dimensional context alongside sequence-based features.

The sequence-level findings further distinguish deletions from insertions. Pathogenic variants occurred at more conserved sites in both types, whereas the associations with indel length and changes in local hydropathy and sequence entropy differed in direction. Pooling deletions and insertions would therefore obscure these associations. Experimental indel maps have similarly shown that the relative tolerance of deletions and insertions varies across proteins and positions [10]. These findings support their separate evaluation and calibration in INDELVAR.

An important consideration is how INDELVAR-derived PP3 and BP4 evidence is combined with PM4 and BP3, respectively. PM4 applies to protein length changes caused by in-frame insertions or deletions in non-repeat regions, as well as stop-loss variants, whereas BP3 applies to in-frame indels in repetitive regions without known function [1]. Earlier guidance advised against combining PM4 with PP3 or BP3 with BP4, while more recent guidance does not impose a general restriction and places greater emphasis on gene- and disease-specific specifications [65–67]. Because INDELVAR includes indel length and repeat-region status among its features, its PP3 and BP4 evidence may not be fully independent of PM4 and BP3. These criteria should therefore be combined according to the applicable specifications.

Several limitations should be considered. Although the test set was separated by release date and contained no training variants, both training and testing relied on ClinVar, and cross-validation was not gene-disjoint. Independent evaluation using variants from other clinical sources is needed. AlphaFold-derived features describe predicted wild-type structures rather than mutant structural consequences and may be less informative in low-confidence or intrinsically disordered regions [68, 69]. INDELVAR was restricted to pure in-frame indels of 1-10 amino acids in MANE Select transcripts, excluding larger or complex events and non-canonical transcripts. Finally, calibration used published pathogenicity priors derived from a broader range of in-frame indels, and the resulting thresholds require further validation in larger independent datasets.

## Conclusions

INDELVAR identifies structural and sequence features that distinguish pathogenic from benign short in-frame indels. By combining these features with gene-level information, the model showed high discrimination in cross-validation and independent testing and reached strong evidence of pathogenicity and benignity for both deletions and insertions upon calibration. The accompanying precomputed resource provides scores and PP3/BP4 evidence for observed GRCh38 in-frame indels.

## List of abbreviations

ACMG/AMP: American College of Medical Genetics and Genomics/Association for Molecular Pathology
AUROC: area under the receiver operating characteristic curve
B/LB: benign/likely benign
CI: confidence interval
ClinGen SVI: Clinical Genome Resource Sequence Variant Interpretation Working Group
DSSP: dictionary of secondary structure of proteins
GERP++: Genomic Evolutionary Rate Profiling
gnomAD: Genome Aggregation Database
IGSR: International Genome Sample Resource
indel: insertion and deletion
INDELVAR: in-frame indel variant pathogenicity predictor
LOEUF: loss-of-function observed/expected upper-bound fraction
LR: likelihood ratio
MANE: Matched Annotation from NCBI and EMBL-EBI
P/LP: pathogenic/likely pathogenic
phyloP: phylogenetic P value
pLDDT: predicted local distance difference test
pLI: probability of loss-of-function intolerance
RSA: relative solvent accessibility
VEP: Variant Effect Predictor
VUS: variant of uncertain significance
WCN: weighted contact number.

## Declarations

### Ethics approval and consent to participate

Not applicable. This study used publicly available variant and protein annotation data and involved no participant recruitment or intervention.

### Consent for publication

Not applicable.

### Availability of data and materials

The data used in this study are available from their original public repositories, as described in the Methods. INDELVAR source code and model are available at https://github.com/eunhui-ji/INDELVAR. The Zenodo deposit provides precomputed scores for publicly observed GRCh38 in-frame indels (https://doi.org/10.5281/zenodo.21285600).

### Authors’ contributions

**Eunhui Ji**: Conceptualization, Methodology, Software, Formal analysis, Investigation, Data curation, Validation, Visualization, Supervision, Project administration, Writing - original draft, Writing - review & editing. **Seung Hwan Oh**: Funding acquisition, Resources, Supervision, and Writing - review & editing. **In-Suk Kim**: Resources and Writing - review & editing. All authors read and approved the final manuscript.

## Acknowledgements

Not applicable.

