## Supplementary Figures for "INDELVAR: structure-informed prediction of in-frame indel pathogenicity with calibrated PP3/BP4 thresholds"

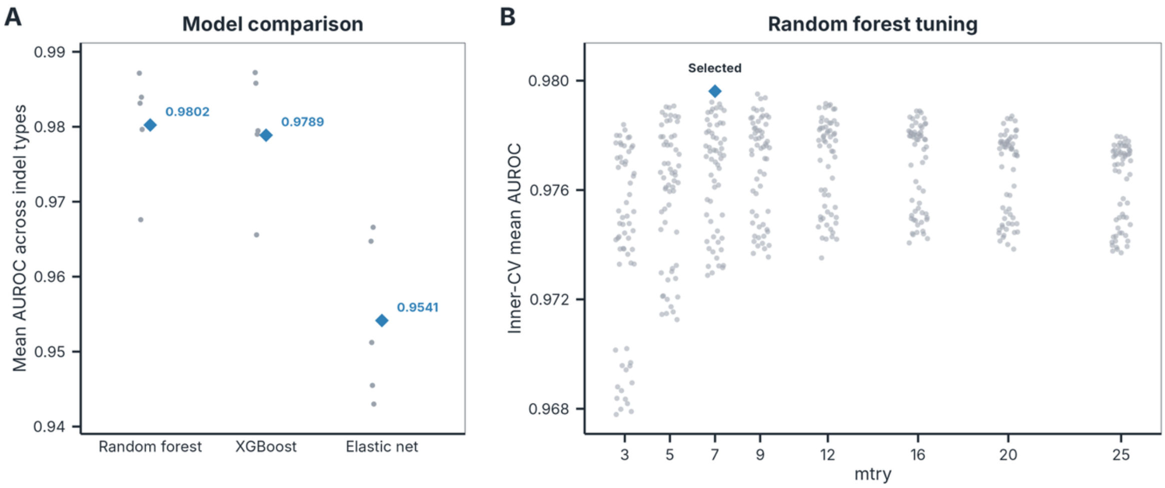


**Supplementary Figure S1**. Model selection and random forest tuning. (A) Comparison of random forest, XGBoost, and elastic-net logistic regression by event-grouped nested cross-validation based on the mean AUROC across indel types; grey points show outer-fold estimates, and diamonds show values from pooled out-of-fold predictions. (B) Inner-cross-validation performance across the complete grid of 480 random-forest configurations; grey points denote configurations and the diamond denotes the selected setting.

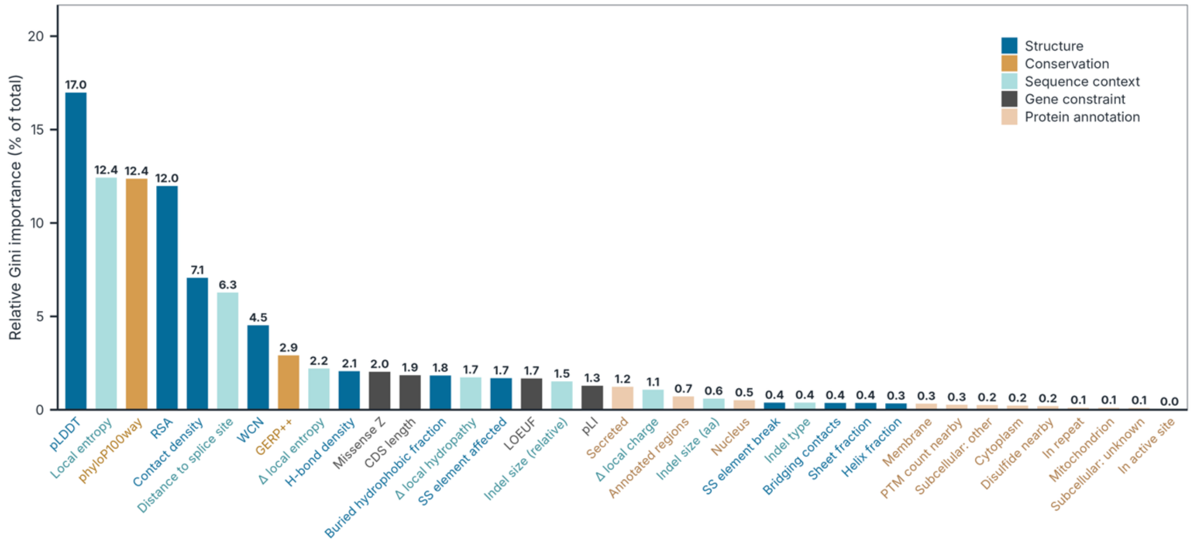


**Supplementary Figure S2**. Gini feature importance in the final INDELVAR model. Relative importance of the 37 model features, calculated from the mean decrease in Gini impurity in the final 1,000-tree random forest and expressed as a percentage of the total. Bars are ordered by importance and colored by feature group.

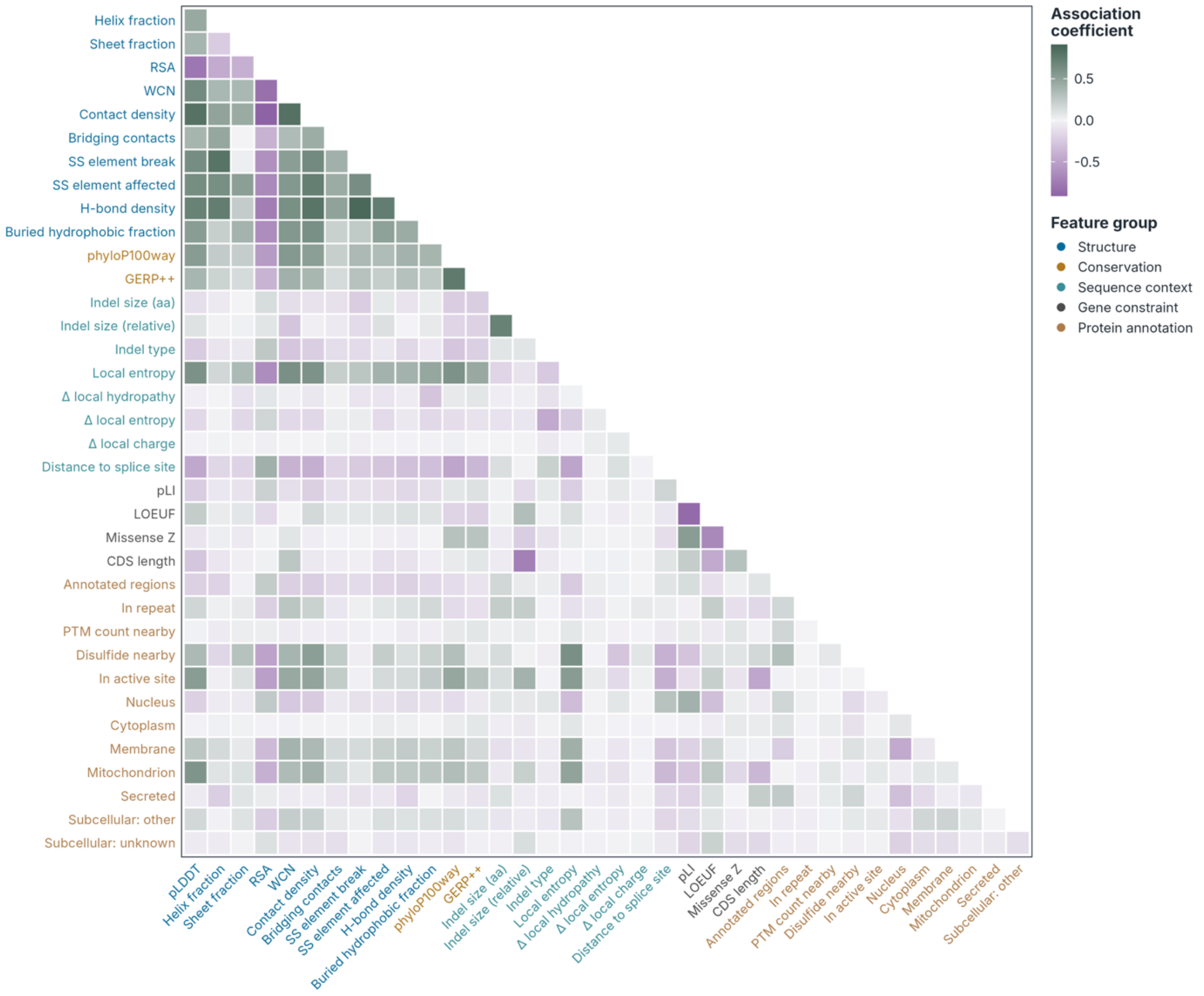


**Supplementary Figure S3**. Pairwise associations among the 37 INDELVAR features. The lower triangle shows associations calculated from pairwise complete observations in the training set. Spearman's rho was used for continuous-continuous pairs, the phi coefficient for binary-binary pairs, and the rank-biserial correlation for continuous-binary pairs. Feature labels are colored by feature group.
